# Beyond competence: a mechanistic model of avian demographic drivers in West Nile virus dynamics

**DOI:** 10.64898/2026.06.15.732287

**Authors:** Elisa Fesce, Eleonora Cattaneo, Giovanni Marini, Roberto Rosà, Davide Lelli, Monica Pierangela Cerioli, Luca Ilahiane, Diego Rubolini, Mario Chiari, Nicola Ferrari

**Affiliations:** Department of Veterinary Medicine and Animal Sciences, Wildlife Health Laboratory, Università degli Studi di Milano, Lodi (LO), Italy; Research and Innovation Centre, Fondazione Edmund Mach, San Michele all’Adige (TN), Italy; Centre Agriculture Food Environment, University of Trento, San Michele all’Adige (TN), Italy; Istituto Zooprofilattico Sperimentale della Lombardia e dell’Emilia Romagna “B. Ubertini”, Brescia (BS), Italy; Department of Environmental Science and Policy, Università degli Studi di Milano, Milano (MI), Italy; Regione Lombardia UO Veterinaria Direzione Generale Welfare, Milano (MI), Italy

**Keywords:** vector-borne infections, mathematical modelling, ecopathology, infection dynamics, avian demography

## Abstract

**Background:** West Nile virus (WNV) is a vector-borne zoonotic pathogen maintained in an enzootic cycle between birds and mosquitoes which is considered a significant public health concern in Europe, particularly in relation to its recent increase in reported human cases and range expansion. While a comprehensive understanding of the virus’s epidemiological dynamics is essential to inform effective prevention and control strategies, to date significant knowledge gaps remain in quantifying interspecific differences within the complex avian communities involved in WNV circulation. Globally, WNV-infection has indeed been documented across more than 300 bird species, however, whether and how inter-specific differences in avian hosts traits affect the spread of WNV is still largely unknown. A substantial body of research has investigated how epidemiological traits, such as the duration of infection and competence, influence WNV dynamics. However, much less is known about the role of avian demography.

**Methodology/Principal findings:** We therefore investigated through mathematical modelling the role of avian demographic traits in shaping patterns of mosquito WNV infection dynamics in northern Italy (Lombardy Region, 2016-2018). We focused on the effects of annual offspring production, timing and synchrony of breeding which ultimately affect seasonal abundance of competent avian hosts. We highlighted that timing of breeding has the greatest effect on the number of infected mosquitoes, while annual offspring production influences the timing of the infection peak. Our simulations provide evidence that non-corvid species can have a key impact on WNV transmission.

**Conclusion/Significance:** These results can support future research by providing priority bird species to direct further studies and by suggesting that the acknowledgment of spatio-temporal variation in the abundance of competent avian hosts plays a key role in the development of effective surveillance strategies and mosquito control actions.

**Author summary:** West Nile virus (WNV) is endemic in Italy and represents a significant public health threat in Europe, with increasing cases of severe neuroinvasive disease in humans in recent years. Surveillance data reveal marked spatial and temporal variability in infection dynamics, suggesting that key drivers of WNV transmission remain poorly understood. The contribution of different bird species (over 300 are implicated in the WNV cycle) is often overlooked despite evidence that species-specific traits are critical determinants of WNV infection dynamics. Few studies have examined birds’ demographic traits, despite their well-established importance in shaping infection dynamics across diseases. Given the challenges in collecting detailed wildlife data, we employed mechanistic models to explore transmission scenarios and test whether avian demographic traits influence bird species’ roles in WNV transmission and maintenance in Lombardy. Our findings demonstrate that brood size, hatching synchrony, and hatching time significantly affect estimated WNV prevalence in mosquitoes.

## Introduction

West Nile virus (WNV) is an emerging zoonotic pathogen belonging to the *Flaviviridae* family. It is maintained in an enzootic cycle involving ornithophilic mosquitoes, primarily belonging to the *Culex* genus [1], and over 300 bird species [2]. Humans and other mammals can be infected through the bite of infectious mosquitoes, but they do not develop sufficient viremia to contribute to virus maintenance and only act as dead-end hosts. In humans, symptoms are rare, but they range from mild flu-like syndrome to severe neuroinvasive disease [3], thus making WNV of public health concern. Furthermore, WNV expansion in new areas [4–6] and the recent observed increase in the incidence of neuroinvasive cases in Europe [4] emphasise the need to better understand the mechanisms driving WNV diffusion, to prevent infections in both humans and animals.

Despite the wide range of bird species considered susceptible to the infection, substantial knowledge gaps continue to hinder the relative contribution of different species to virus maintenance and transmission. Host competence depends on several factors such as the duration of infection (shaped by immune clearance or host mortality), the intensity of viraemia, infection lethality, contact or biting rates, and host abundance and density, all subordinated to susceptibility to the infection. Most of these factors are challenging to assess in natural settings, are highly species-specific [7–10], and often remain largely overlooked and poorly quantified especially for European bird species [7–13]. The species-specific contributions to WNV dynamics can therefore vary, and the spatial and temporal heterogeneity in avian host population size and community composition may play a role in WNV transmission [14]. In addition to that, demographic traits of host species are known to shape infection dynamics. For instance, two key metrics of infection dynamics, the basic and effective reproduction numbers (*R₀*and *R*, respectively), are strongly influenced by the abundance of susceptible individuals and their natural mortality. These numbers represent the expected number of secondary infections generated by a single infectious individual, respectively in a fully susceptible population (*R₀*) or over time (*R*), thereby highlighting the role of host abundance and demographic traits in shaping infection dynamics [15–17]. Furthermore, host demographic factors such as fecundity and seasonal patterns in breeding time affect the number and proportion of susceptible individuals (newborns and fledglings are usually susceptible to infections, either from birth or once maternal immunity wanes a few months later), thereby modulating pathogen transmission [18–23]. Also, demography influences local population size, thereby determining whether the infection might fade out or persist over time [24,25]. Together, these demographic aspects contribute to pathogen dynamics but are yet often overlooked in epidemiological studies in wildlife populations, where monitoring and surveillance are more challenging [26].

Although data on bird species occurrence and relative abundance from ornithological surveys are potentially available, as are typical ranges of reproductive parameters (e.g. breeding season, number of broods, and clutch size), relatively few studies have explicitly incorporated these factors when investigating the dynamics of WNV [27]. Moreover, the high number of species involved in WNV transmission, together with the complexity of conducting extensive field and laboratory investigations, makes it difficult and resource-intensive to determine the specific role of each host in natural settings.

Analytical and theoretical approaches can complement field studies by investigating alternative transmission scenarios and testing hypotheses that can then be verified in the field. While to date a substantial branch of disease ecology modelling relies on statistical approaches starting from data and inferring patterns from observations, mechanistic or mathematical models offer a complementary perspective: they allow both the evaluation of mechanisms of transmission by calibrating models on real data and allow the exploration of ‘what-if’ scenarios. By systematically varying transmission assumptions or specific parameters, mathematical models can generate theoretical predictions about alternative conditions thus providing a framework to formulate hypotheses on how the system might behave under different assumptions.

We employed a mathematical model calibrated to reproduce WNV dynamics in the Lombardy Region, an endemic area for this pathogen in northern Italy, to generate ‘what-if’ scenarios under different demographic conditions of bird demography. This approach aimed to evaluate whether variations in avian host breeding traits could influence the seasonal dynamics of WNV transmission. We used a temporally explicit mathematical model to explore the effects of three main avian demographic traits (mean annual fledgling production, synchrony of clutches and timing of hatching) on the seasonal dynamics of mosquito-borne WNV infection. Our research relied on a previously validated framework that simulates WNV dynamics in birds and mosquitoes [28–30] that uses data collected during regional WNV surveillance campaigns [31] in Lombardy Region, across multiple sites. The model includes a time-dependent function to simulate seasonal reproduction cycles [32], which accounts for annual offspring production, the breeding synchrony of the timing of breeding cycles, and for the variable timing of breeding. These three variables were varied within plausible biological ranges, extrapolated from the literature, to theoretically investigate their impact on the temporal daily variation in the number of WNV-infectious mosquitoes.

## Methods

Our study is structured into three interconnected components addressing the effects on seasonal WNV dynamics of avian demographic traits (Fig 1):

**Fig 1.**
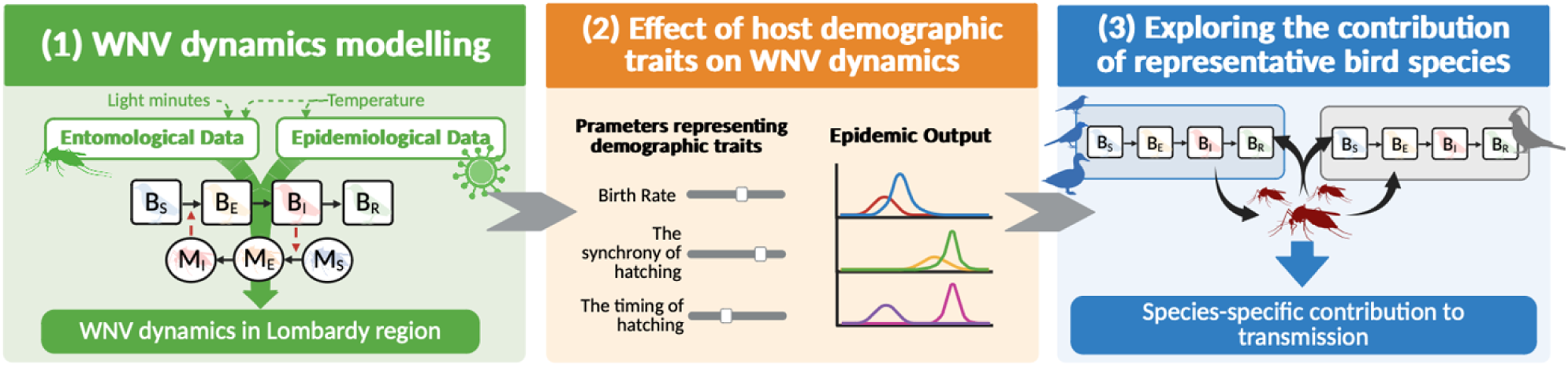
Schematic representation of the study. (1) Calibration of the baseline WNV transmission model using entomological and surveillance data to obtain a reliable framework to simulate WNV dynamics in Lombardy Region. (2) Exploration of the ‘what-if’ scenarios through alterations of birds’ demographic parameters, assessing potential impacts on transmission dynamics. (3) Extension of the baseline model to include two avian host species, enabling quantification of species-specific contributions to WNV transmission. Created in BioRender. Fesce, E. (2026) https://BioRender.com/z9etma0.

### 1. WNV dynamics modelling

First, building on the model proposed in [28], we simulated WNV dynamics across the Lombardy Region over the study period (2016-2018). Given the complexity of the system and the fact that several mechanisms are still unknown or unquantified, this step was crucial to estimate the unknown (or location-specific) parameters and provide an accurate simulation of WNV dynamics in the study area. The temperature-dependent model was fitted to entomological and WNV surveillance data from the Italian national arbovirus surveillance plan gathered in the region, following the methodology proposed in the original study [28], and provided the baseline model for the following steps.

### 2. Effect of host demographic traits on WNV dynamics

The calibrated model was used to explore a set of ‘what-if’ scenarios, focusing on variation in host demographic traits, with the aim of assessing their potential influence on transmission dynamics under alternative assumptions.

### 3. Exploring the contribution of representative bird species

Finally, the baseline model was further extended to explicitly include two avian host species, allowing us to investigate and quantify the contribution of representative existing species to WNV abundance in mosquitoes.

### WNV dynamics modelling

#### Modelling framework description

Our modelling framework consists of two deterministic compartmental model mimicking the mosquito (*Culex pipiens*) lifecycle (entomological model) and the WNV transmission dynamics between the vector and host populations (epidemiological model). Specifically, the latter translates biological assumptions on WNV transmission into a system of differential equations representing the underlying infection processes. It describes the temporal dynamics of a competent bird species and a mosquito population, both structured by epidemiological status, and explicitly accounts for host–vector interactions and temperature influence on mosquito survival and infection mechanisms. The framework was originally developed [29,30] for a single competent bird species (*Pica pica*, magpie) but can be readily adapted to represent other species by including species-specific demographic and epidemiological parameters. In this study, however, all avian host trait parameters (e.g. birth rate) were estimated by fitting the model to mosquito epidemiological data, starting from a plausible abundance of birds within the mosquitoes’ home range (S1 Text). Therefore, the simulated bird population represents a generic competent host rather than any specific species. This choice was motivated by the limited epidemiological WNV-related information available for most European bird species regarding their role in WNV transmission. Such knowledge gaps constrain our ability to identify precise reservoir species in Europe and, combined with the ecological complexity of avian communities, justify our decision to model a general competent bird reservoir rather than focusing on a single species.

Host and vector populations are divided into compartments according to their infection status (susceptible, infected, and infectious), enabling the estimation of WNV circulation dynamics. The model structure follows that of the original study [28], with modifications to incorporate avian demographic dynamics. Following [32], time-dependent periodic cosine function representing the daily per capita offspring production was integrated allowing a more realistic demography in birds. We refer to this function as the birth function and according to the original study it depends on three parameters, the annual per capita birth rate (here the annual offspring production), the timing of the peak of the birth pulse and the duration of the birth pulse. Avian compartments were simplified to four (representing susceptible, exposed, infectious and recovered birds, with no age class subdivision) due to limited information on the local age structure and the absence of precise juvenile parameters. These adjustments allow the model to focus on the impact of avian demographic traits on WNV dynamics while preserving the core structure and assumptions of the original framework. The model simulates infection dynamics between birds and mosquitoes, based on observed adult female mosquito abundance and WNV prevalence (*Culex* genus, grey box in Fig 2). Briefly, WNV transmission occurs exclusively through mosquito bites, with the transmission rate determined by the number of infectious birds and mosquitoes, the average number of mosquito bites per day, and the proportion of bites directed at birds. Birds can recover from infection and develop lifelong immunity (SEIR-type structure, Fig 2), whereas infected mosquitoes remain infectious for their entire lifespan, given their short life expectancy (SEI model, Fig 2). The framework incorporates an incubation period in both hosts and vectors. Full model details, including equations and parameter values, are provided in S2 Text, while a schematic representation of the model structure is shown in Fig 2. The daily number of mosquitoes is first estimated through the entomological sub-model (represented by the grey box in Fig 2), which provides the abundance of circulating adult female mosquitoes, whereas bird dynamics are explicitly modelled through the time-dependent birth function *b(t)*.

**Fig 2.**
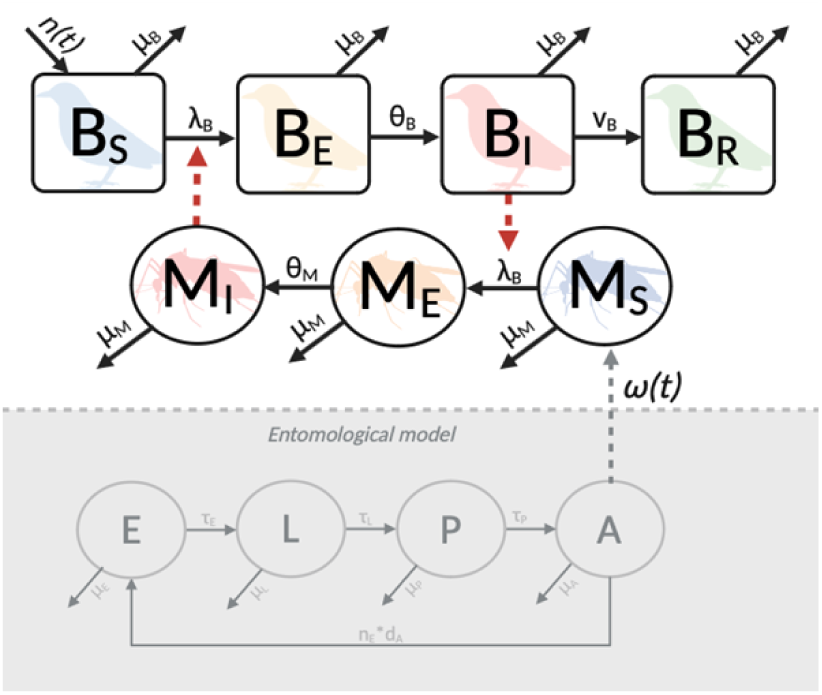
Model scheme. Schematic representation of the modelling framework. Squares represent the compartments describing the bird population: B_S_, B_E_, B_I_ and B_R_ indicate the susceptible, exposed, infectious, and recovered birds, respectively. Circles represent the mosquito population (*Culex pipiens*), with M_S_, M_E_ and M_I_ corresponding to susceptible, exposed, and infectious mosquitoes. Red dashed arrows indicate the transmission rates between compartments. The entomological sub-model estimating the daily number of mosquitoes is represented by the grey box, where circles represent the four developmental stages of mosquitoes: E for eggs, L for larvae, P for pupae, and A for adults. Created in BioRender. Fesce, E. (2026) https://BioRender.com/z9etma0.

#### Model parameters

Where available, model parameters were derived from the literature (see Table 1 in S2 Text for known parameters). Parameters that could not be obtained from published sources were inferred using a Markov Chain Monte Carlo (MCMC) approach, as described in Fesce et al. [28] (see Table 2 in S2 Text for parameters estimated via MCMC and description of the method). This procedure allowed us to identify the range of parameter values most consistent with the observed mosquito abundance and WNV prevalence. The mean parameter values obtained from this re-estimation were then used to specifically assess the influence of avian demographic traits on WNV dynamics. In accordance with the original model and previous studies showing that mosquito developmental rates and the probability of WNV transmission from birds to mosquitoes are influenced by temperature [33–38], the modelling framework incorporated mean daily temperature data recorded by ARPA Lombardia (https://www.arpalombardia.it), following the methodology described by [28–30]. Biological interpretations of all model parameters are provided in the S1 Text.

**Table 1.** Avian demography parameters description and biological meaning.

| Parameter name | Description | Derivation criteria |
| --- | --- | --- |
| Birth rate ($\mu_B$) | Mean annual offspring production | $\mu_B$ accounts for the offspring production per clutch and the mean number of clutches per year* |
| Synchrony of hatching ($s$) | Bandwidth of the birth function, i.e. the duration of the reproductive season (offspring production) | $s$ was chosen to yield 90% of the annual recruitment output within the bandwidth. Parameter value was constrained to produce an overall timing compatible with published phenology and expert opinion** |
| Timing of hatching ($\varphi$) | $\varphi$ determines when hatching reaches its seasonal peak. | $\varphi$ was calibrated so that the day at which the function attains its maximum falls near the midpoint of the species' breeding season, in agreement with available literature and expert opinion** |

\* Values estimated from literature
\*\* In the absence of detailed quantitative data on the timing of peak hatching and the temporal distribution of births, parameter values were chosen to yield recruitment curves consistent with the known reproductive phenology of each species.

**Table 2.** Ranges of parameters for the sensitivity analysis.

| Parameter | Biological interpretation of parameters | Baseline value* | Range** |
| --- | --- | --- | --- |
| $\mu_B$ | Mean birth rate | 0.0079 | 0 - 0.022<br>(0 – 8 offspring/year) |
| $s$ | Synchrony of hatching | 11.84 | 5-20 |
| $\varphi$ | Timing of hatching | 0.59 | 0.33- 0.66<br>(April 30<sup>th</sup> – September 1<sup>st</sup>) |

\* The baseline value was obtained from point (1) *WNV Dynamics Modelling*, by averaging the values obtained by fitting the model on Lombardy Region data.
\*\* Plausible intervals considering the breeding biology of bird species that occur in Lombardy Region. Ten equally spaced values were sampled across each interval for each parameter.

#### Dataset and model calibration

Entomological and epidemiological dataset: Temporally explicit models of WNV dynamics were calibrated on the entomological surveillance data collected in the Lombardy Region within the national surveillance plan for arboviruses between 2016 and 2018 [31]. Mosquito abundance records informed the calibration of unknown parameters in an initial entomological dynamic model, which was designed to estimate daily mean mosquito abundance. Subsequently, WNV pool-positivity data were employed to estimate the epidemiological parameters governing WNV transmission across the study region. The 40 surveillance sites were grouped to estimate the average daily mosquito abundance and WNV dynamics in Lombardy Region.

### Effect of host demographic traits on WNV dynamics

To investigate whether changes in avian demographic traits affect WNV dynamics, we modified key demographic parameters values of the avian hosts in the baseline model, one at a time, following a sensitivity-analysis–type approach. We then evaluated the resulting effects on the estimated number of infectious mosquitoes relative to baseline simulations. This procedure allowed us to isolate the influence of individual demographic traits on transmission dynamics. Specifically, we focused on the three parameters that are likely to differ across species and to play a relevant role in WNV circulation and that characterize the birth function: 1) the birth rate (*µ_B_*), 2) the synchrony of hatching (*s*) and 3) the timing of hatching (*φ*). Parameter explanation and biological meaning are reported in Table 1.

For the analysis, ten equally distanced parameters values for demographic traits were chosen within biologically plausible ranges that could be representative of most bird species breeding in the study area, as defined through a targeted literature review (details of the review and sources are in the section “Bird traits”). Each theoretical scenario explored the effect of a variation of the reproductive demographic traits of the avian population, representing the diversity observed among competent bird species in the Lombardy Region.

Starting from the baseline model, parameterised with field-derived estimates, each trait was independently modified while keeping the others constant, to isolate its impact on transmission dynamics throughout the mosquito activity season. All simulations were performed under identical environmental and entomological conditions, i.e. using the same underlying temperature, daylight and daily mosquito abundance. Demographic parameters were varied within their biologically plausible range (Table 2, and the resulting birth functions are reported in Fig S2 in S2 Text), while the remaining parameters were held constant at their initial estimates, corresponding to the mean values derived from baseline simulations across the study area. The use of averages decreases the realism of the simulations, but makes it possible to compare the effects of changing individual parameters, thus allowing us to focus on the effects of investigated parameters. The outcomes were then compared to the baseline scenario to quantify how species-specific characteristics modulate infection dynamics.

### Exploring the contribution of representative bird-species

Finally, we extended the analyses by adapting the model structure to explore the specific contribution of WNV-competent bird species. As a first step, we selected the bird species to include in the model that were both competent for WNV and differed in key demographic traits. We then incorporated their species-specific traits into the modelling framework. This approach allowed us to evaluate whether, and to what extent, realistic demographic characteristics influence WNV transmission dynamics.

#### Selection of bird species

To select the species to be included in our modelling framework, we focused on birds that have been shown to be capable of transmitting the virus [2,7–10]. From this subset, species were further selected based on the following criteria:

1. Breeding in the Lombardy Region [39], in order to align avian and mosquito data that were collected exclusively in this region and during the peak of mosquito activity.
2. Breeding in lowland temperate areas, to ensure ecological consistency with typical WNV-competent mosquito habitats.
3. Inclusion in the National Plan for Prevention, Surveillance, and Response to Arboviruses 2020–2025 [40], which lists species of epidemiological relevance.
4. Mosquito feeding preference, based on data from Zeller et al. [41] only species that are preferentially targeted by mosquitoes were included.

Among the potentially competent bird species for WNV transmission, the following 13 bird species meet our inclusion criteria: *Anas platyrhynchos*, *Ardea cinerea*, *Columba palumbus*, *Corvus cornix*, *Dendrocopos major*, *Garrulus glandarius*, *Parus major*, *Passer montanus*, *Pica pica*, *Streptopelia decaocto*, *Sylvia atricapilla*, *Turdus merula*, *Vanellus vanellus*.

#### Clustering of bird species for WNV dynamics analyses

To reduce the analysis to species representative of different demographic profiles, we performed a cluster analysis to identify groups of species sharing similar demographic traits. The species-specific traits for all 13 selected taxa included in the cluster analysis were compiled as described in the following section “Bird traits” and are provided in Table 2 in S1 Text.

A k-means clustering was performed on the selected species using the cluster package in R (version 4.5.1). The identification of the optimal number of clusters was based on the highest silhouette coefficient. The clustering incorporated the following characteristics:

- Relative abundance of the species in the Lombardy Region
- Annual Growth Rate (Annual proportional change in population size)
- Breeding season (start and end of the estimated period of reproductive activity based on species-specific phenology)
- Timing of egg laying
- Number of clutches laid/offspring raised per pair per year
- Clutch size
- Incubation period duration
- Hatching success
- Fledgling period duration
- Age at first breeding
- Average lifespan
- Mortality rates for nestlings, juveniles and adults

All parameter values included, and their relative sources are reported in detail in S1 Text.

#### Bird traits

Bird parameters values representing bird demographic traits were obtained from published sources. We conducted a literature review using PubMed and Google Scholar, searching for scientific articles that included the scientific names of the bird species combined with keywords related to the parameters of interest (detailed description of the search strategy and key words used can be found in S1 Text). To supplement the information retrieved from the literature, we also incorporated data from the British Trust for Ornithology (BTO) [42] which offers comprehensive coverage of our species of interest. This resource was instrumental as it provides standardized metrics across target species, ensuring methodological consistency and facilitating robust comparison among them. Where multiple sources were available, we prioritized the most frequently reported values or those from regions with similar microclimatic conditions to Lombardy region. Furthermore, whenever possible, we selected parameters from the same source across different species to ensure internal consistency. The list of all included parameters, their biological meaning and the complete database including all parameters associated with each bird species are provided in S1 Text.

#### Species contribution to WNV spread

Subsequently, three species were selected as representative of the identified clusters: *Turdus merula, Pica pica* and *Anas platyrhynchos.* According with literature information (S1 Text), simulation parameters of the three species are reported in Table 3.

**Table 3.** Species specific demographic ed epidemiological parameters for WNV of the three representative avian species selected by the cluster analysis.

| Bird species | Mosquito biting rate on the species ($b_1$)* | Mean birth rate ($\mu_B$)** | Synchrony of hatching (s)** | Timing of hatching ($\varphi$)** | Recovery rate ($\nu_B$)*** | Incubation period ($\theta_B$)*** |
| --- | --- | --- | --- | --- | --- | --- |
| *Turdus merula* (blackbird) | 0.35 | 0.013<br>(4.8 offspring/ year/ individual) | 6<br>March to June | 0.35<br>8 May | 1 | 0.22 |
| *Pica pica* (magpie) | 0.1 | 0.0074<br>(2.7 offspring/ year/ individual) | 6<br>April to July | 0.4<br>26 May | 0.66 | 0.2 |
| *Anas platyrhynchos* (mallard) | 0.01 | 0.014<br>(4.2 offspring/ year/ individual) | 10<br>mid-March to mid-June | 0.33<br>30 April | 0.66 | 0.25 |

\*From Rizzoli 2015 [43], unpublished data
\*\*Details and references of the estimated birth functions resulting from parameters in Table 3 are reported in Fig S3 in S2 Text.
\*\*\*Epidemiological values were derived from [8]. The incubation period was estimated as the inverse of the duration of viremia.

It should be noted that the clustering was intended solely to simplify subsequent modelling analyses and has no taxonomic or ecological significance. The species selected as representative of the clusters were chosen among species widespread and abundant within the study area and primarily based on the availability of epidemiological data for WNV and their confirmed involvement in the enzootic cycle of the infection (e.g. duration of viremia in the species).

The contribution of the three representative species to WNV dynamics was estimated by modifying the model to include two competent bird species: one known species, which represent in turn one of the three species selected (for specific parameter values see Table 2), and a second generic species, that is competent and for which parameters were estimated from epidemiological data via MCMC as in the baseline model. In detail the model was modified by including further four additional differential equations to represent the selected species (equations reported in S2 Text), resulting in a 11 differential equations model (model scheme reported in Fig 3). The selected species were incorporated individually, one at a time, and for each simulation, the model was rerun using entomological data for the region. Parameters for the second bird population, representing the remainder of the avian community, were then estimated using MCMC, allowing the model to account for interactions between the selected species, the overall bird population, and mosquitoes. This resulted in an extended model simulating WNV dynamics between mosquitoes and two bird populations, one representing the chosen species and the other representing the rest of the avian community, [44]. This approach enabled us to evaluate whether, and to what extent, realistic species-specific demographic characteristics affect WNV transmission dynamics.

**Fig 3.**
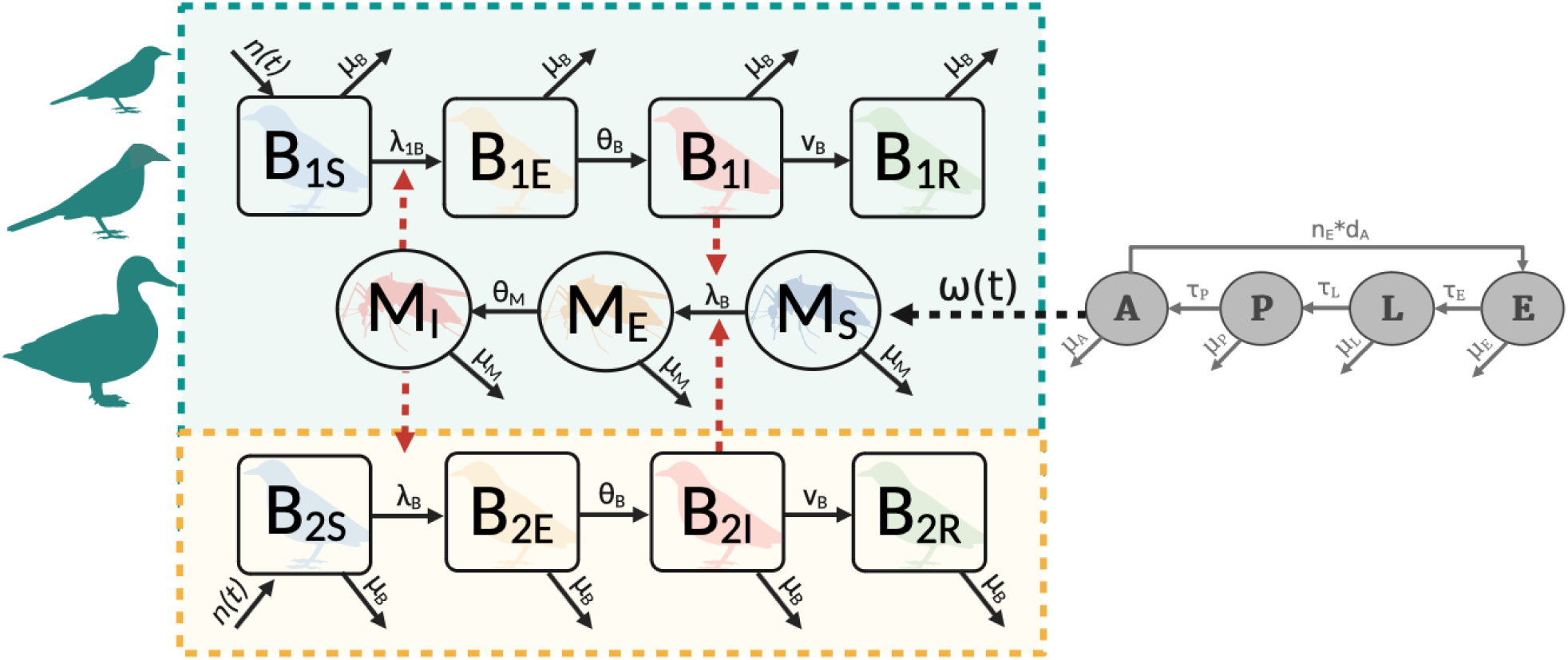
Model scheme for the modified modelling framework. Squares represent the bird population, B_S_, B_E_, B_I_ and B_R_ for susceptible, exposed, infectious and recovered birds respectively. Birds are represented in two compartments: B_1_ the cluster-selected species (green shaded area), B_2_ the rest of avian community (orange shaded area). Circles represent mosquitoes, with M_S_, M_E_ and M_I_ representing the number of susceptible, exposed and infectious mosquitoes (*Cx. pipiens*) respectively. The grey circles represent the entomological model underlying the estimates of the number of circulating adult female mosquitoes (ω(t)). The green shaded area of the figure denotes the model components informed by species-specific parameters derived from the literature review, whereas the orange shaded area indicates the parameters estimated by the model through the MCMC process. Created in BioRender. Fesce, E. (2026) https://BioRender.com/z9etma0.

The specific contribution of each species on WNV transmission was assessed by calculating the number of new infected mosquitoes resulting from mosquito bites on that bird species, relative to the total number of new infectious mosquitoes:

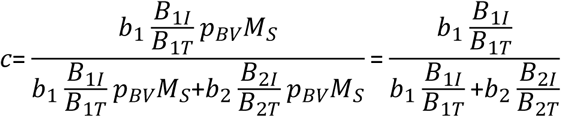

Where *c* represents the specific contribution of each species; *b_1_* is the specie specific biting rate for the selected bird species population; *B_1I_*is the daily number of infected birds of the known population; *B_1T_*is the total number of birds of the known population; *b_2_* is the biting rate for the competent avian community (*B_2_*); *B_2I_*is the daily number of infected birds of the competent avian community; *B_2T_*is the daily total number of the competent avian community; *p_BV_*is the probability of WNV transmission from bird to mosquito per infectious bite and *M_S_* is the daily number of susceptible mosquitoes. We therefore expressed the contribution of each species as a value between 0 and 1, where 0 indicates that no new infections can be attributed to the investigated species, and 1 indicates that 100% of new infections are attributable to it.

## Results

### WNV dynamics modelling

The base model describing WNV dynamics in Lombardy Region provided consistent estimates with the observed viral circulation in mosquitoes, showing an increase in the number of infectious mosquitoes in July, peaking in August/September, and declining in October. The predicted number of infectious mosquitoes was highest in 2018, followed by 2017 and lower in 2016. In all the three years of simulation the trend of the number of positive samples is well captured by the model, and the 75% of observations (red dots in Fig 4) fall within the 25^th^ and 75^th^ percentiles of predictions (blue boxes Fig 4).

**Fig 4.**
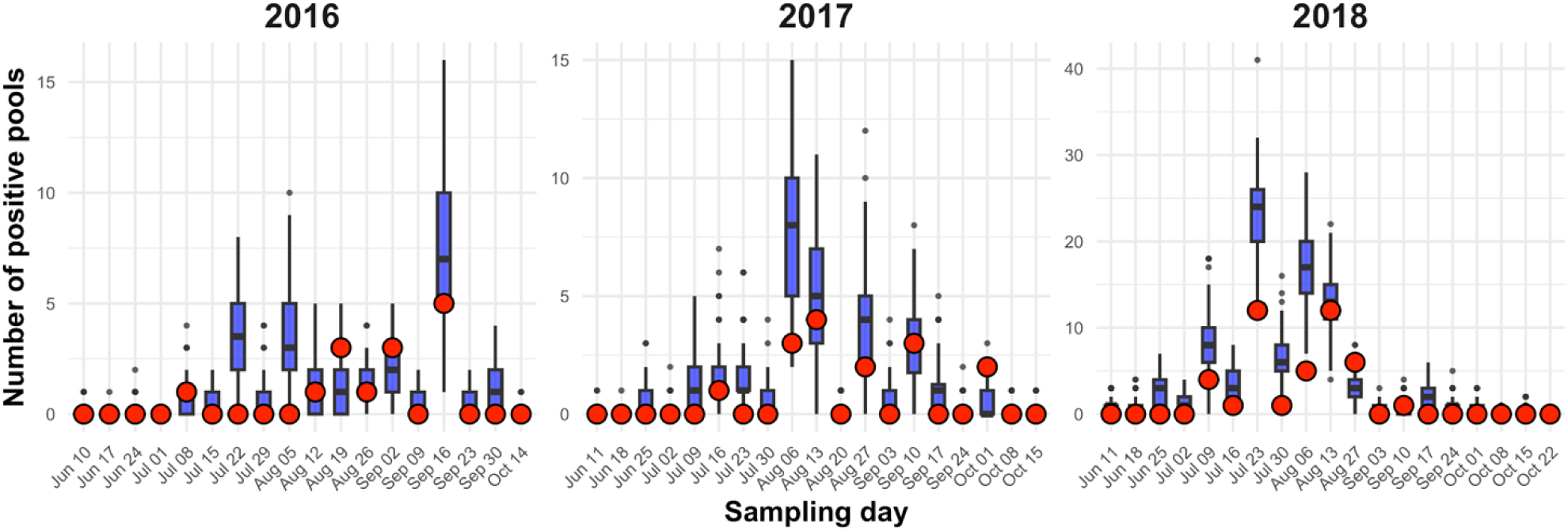
Base model fit. Yearly fit of the model, with red dots representing the observations and blue boxplots showing the distribution of predicted WNV-positive pools. The lower and upper hinges of the boxplot represent the 25^th^ and 75^th^ percentiles, the central line denotes the median, and the whiskers extend to the most extreme values within 1.5 times the interquartile range (IQR). The axes are scaled differently across years to align with year-specific sampling periods and to optimise data visualisation.

The model parameter estimation via MCMC yielded a birth function that varied across years, reflecting interannual variability in bird reproductive dynamics, with the maximum mean daily number of offspring reaching 0.042 in late August 2016, 0.021 at the end of July 2017, and 0.069 in late September 2018 with the hatching season consistently lasting approximately four months (Fig 5). From visual inspection, the overall synchrony of hatching does not appear to differ substantially among years. Estimates for all free parameters are presented in S2 Text.

**Fig 5.**
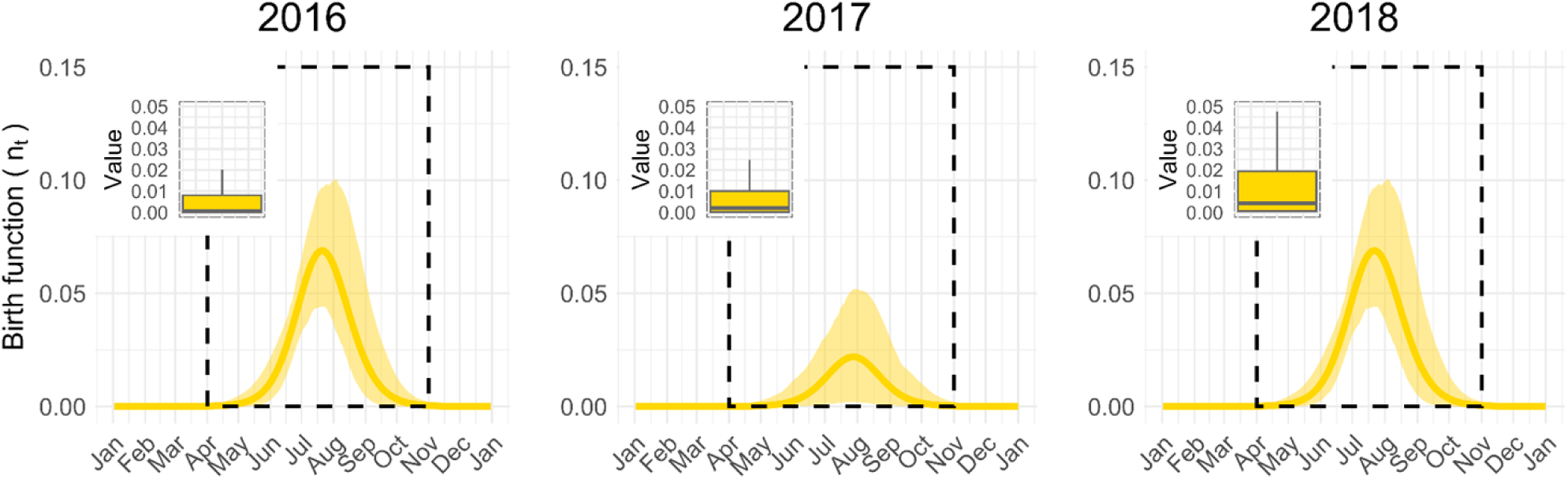
Predicted birth function. Mean predicted birth function (yellow solid line) with 95% confidence intervals (yellow shaded area) for 2016, 2017, and 2018. The dashed black box indicates the mosquito activity period. The gold boxplot shows the annual distribution of predicted values across simulations (mean ± 95% CI).

### Effects of demographic traits on WNV dynamics

#### Effect of the number of newborn (*µ_B_*)

As number of newborn birds increases (higher *µ_B_*), the peak in the number of infectious mosquitoes advanced from August to July. The impact of the number of newborns (growth rate *µ_B_*), however, varies across the three simulated years. In 2016 and 2017, years characterised by milder WNV circulation and fewer infectious mosquitoes (see Fig 6A), a reduction in the number of newborns (*µ_B_*) leads to both an increase in the number of infectious mosquitoes and a delay in the peak, shifting it towards late summer. In contrast, in 2018 a reduction in the number of newborns (*µ_B_*) results only in a temporal shift of the peak, without a substantial increase in the number of infected mosquitoes (Fig 6A).

**Fig 6.**
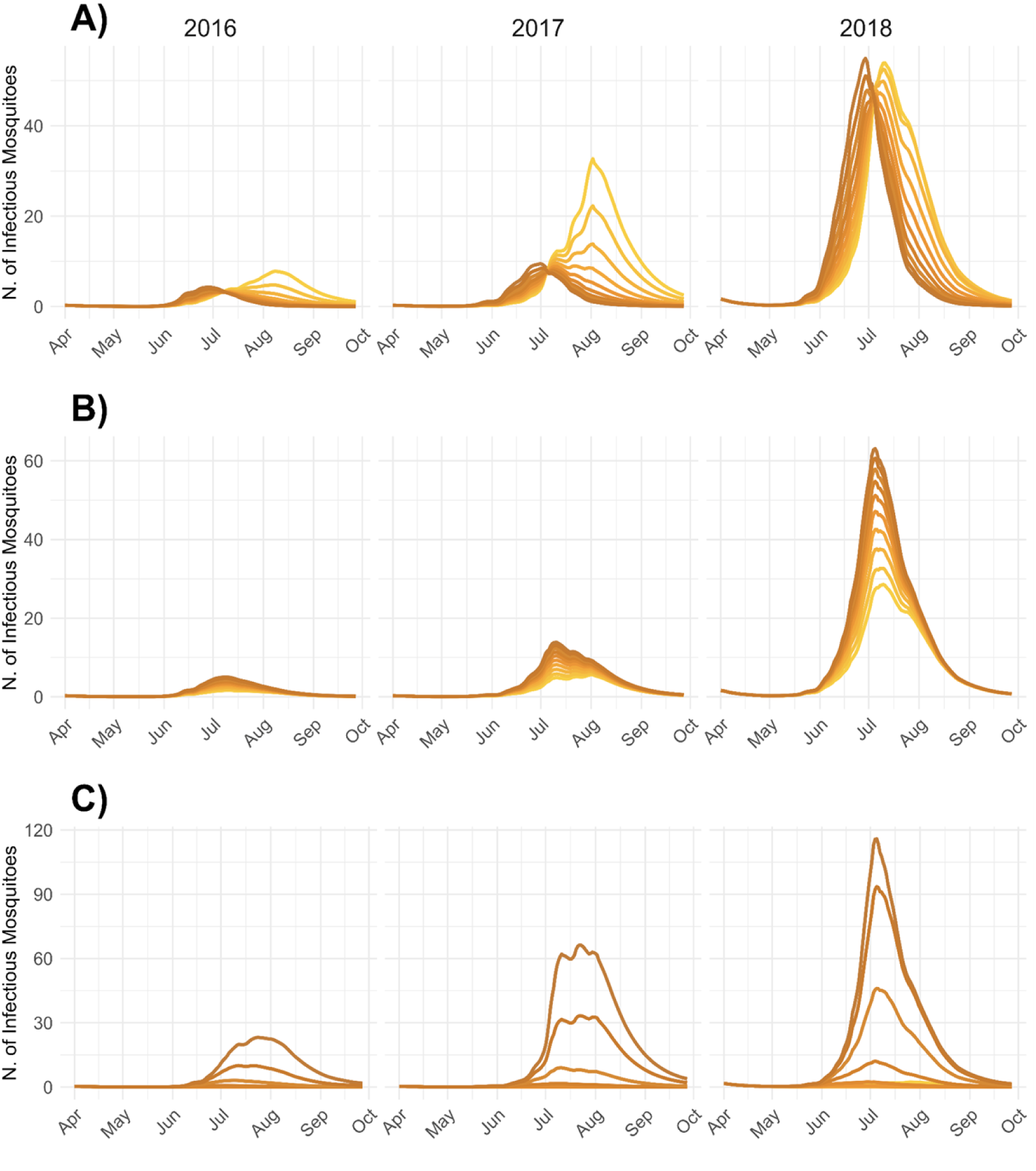
Effect of the demographic parameters. (A) Predicted average number of infectious mosquitoes as the number of newborn birds (*µ_B_*) increases across three years of simulation. The y-axis represents the number of infectious mosquitoes, with darker curves indicating higher numbers of newborn birds (values of *µ_B_*: 0, 0.0024, 0.0048, 0.0073, 0.0097, 0.0122, 0.0146, 0.0170, 0.0195, 0.0219, values estimated). (B) Predicted number of infectious mosquitoes as the synchrony of hatching (*s*) increases across three years of simulation. The y-axis represents the number of infectious mosquitoes, with darker curves indicating higher synchrony (values of s: 5, 6.6, 8.3, 10, 11.6, 13.3, 15, 16.6, 18.3, 20). (C) Predicted number of infectious mosquitoes as the timing of hatching shifts *φ* from May to October. The y-axis represents the number of infectious mosquitoes, with lighter curves corresponding to earlier hatching timings and darker curves indicating later ones. The tested values of *φ* (0.33, 0.36, 0.40, 0.47, 0.51, 0.55, 0.58, 0.62, 0.66) correspond to hatching peaks occurring on the following dates: April 30^th^, May 10^th^, May 24^th^, June 19^th^, July 4^th^, July 19^th^, July 30^th^, August 13^th^, and August 28^th^, respectively. Hatching timings earlier than June 19^th^ result in fewer than 15 infectious mosquitoes and are therefore less distinguishable in the figure. The y-axis scales differ among panels to improve the visibility of the plotted patterns.

#### Effect of the synchrony of hatching (*s*)

The synchrony of hatching (*s*) influences the number of infectious mosquitoes but not the timing of the peak. As the hatching season becomes more spread during the mosquito activity period (lower values of *s*), the number of infectious mosquitoes decreases, and vice versa (Fig 6B).

#### Effect of the timing of hatching (*φ*)

The timing of hatching (*φ*) significantly influences the number of infectious mosquitoes, with an earlier birth season leading to a lower peak in the number of infectious mosquitoes (Fig 6C).

We therefore found that all three tested demographic parameters influence WNV dynamics, with the timing of hatching (Fig 6C) having the greatest effect on the number of infectious mosquitoes, and the number of newborns (Fig 6A) influencing the timing of the infection peak.

### Species contribution

#### Cluster analysis of bird demographic traits

According to the cluster analysis, three groups of birds were identified (Fig 7):

- Cluster 1: *Streptopelia decaocto*, *Columba palumbus*, *Turdus merula*, *Passer montanus*, *Sylvia atricapilla*
- Cluster 2: *Corvus cornix*, *Garrulus glandarius*, *Pica pica*, *Dendrocopos major*, *Parus major*
- Cluster 3: *Ardea cinerea*, *Vanellus vanellus*, *Anas platyrhynchos*.

**Fig 7.**
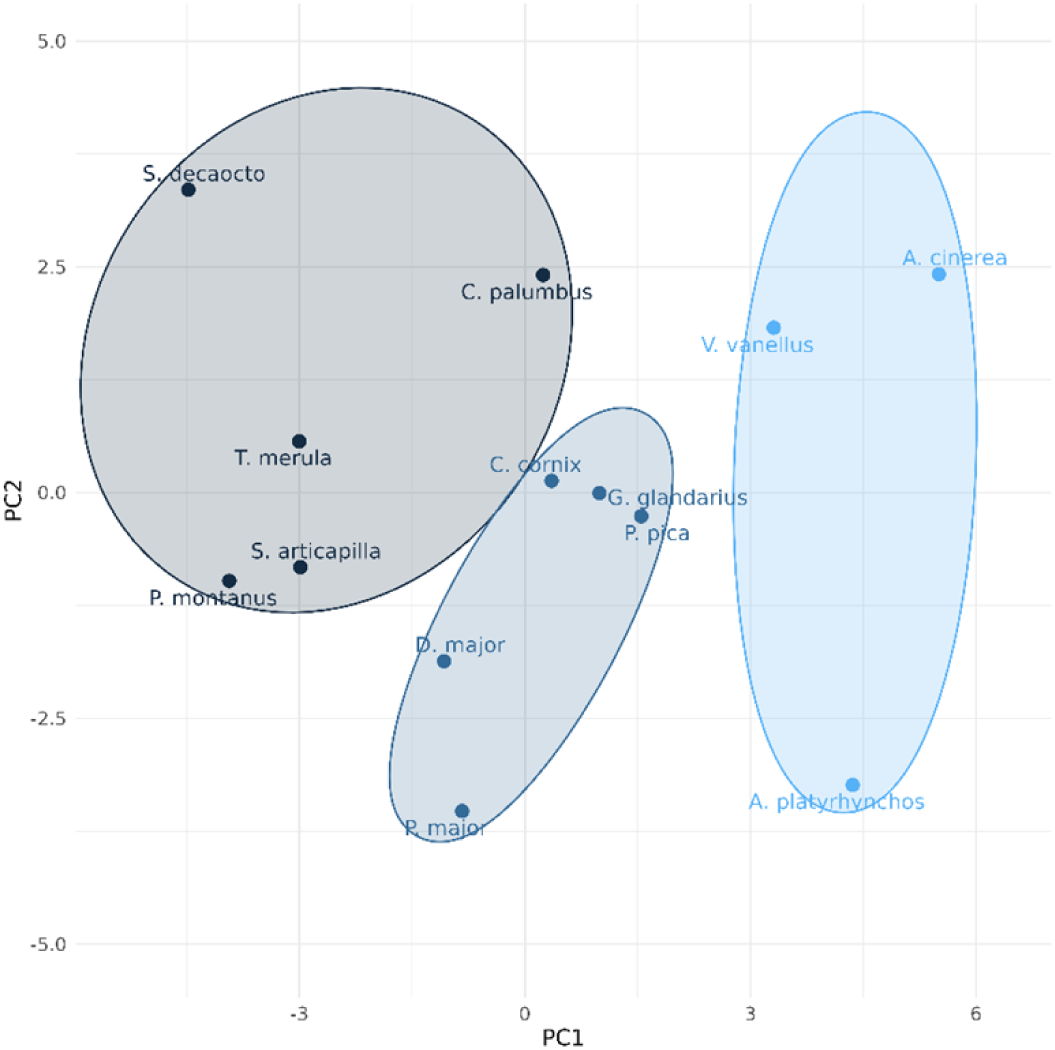
Clustering of birds based on demographic traits. The black, dark blue, and light blue shaded circles represent the three clusters identified through the k-means clustering analysis. Smaller dots, labelled with species scientific names, represent single bird species.

#### Species contribution to WNV spread

Simulations showed that all three representative species can contribute to WNV spread among mosquitoes (Fig 8). The model fit remains robust across the three species, with over 90% of observations falling within the 95% confidence interval of the simulations. Additionally, 71.4%, 73.2%, and 71.4% of observed mosquito prevalence fall within the interquartile range (25^th^–75^th^ percentile) of the simulations assuming magpie, blackbird, and mallard as avian species in the model respectively, as shown by the green, orange and purple boxplots in Fig 8.

**Fig 8.**
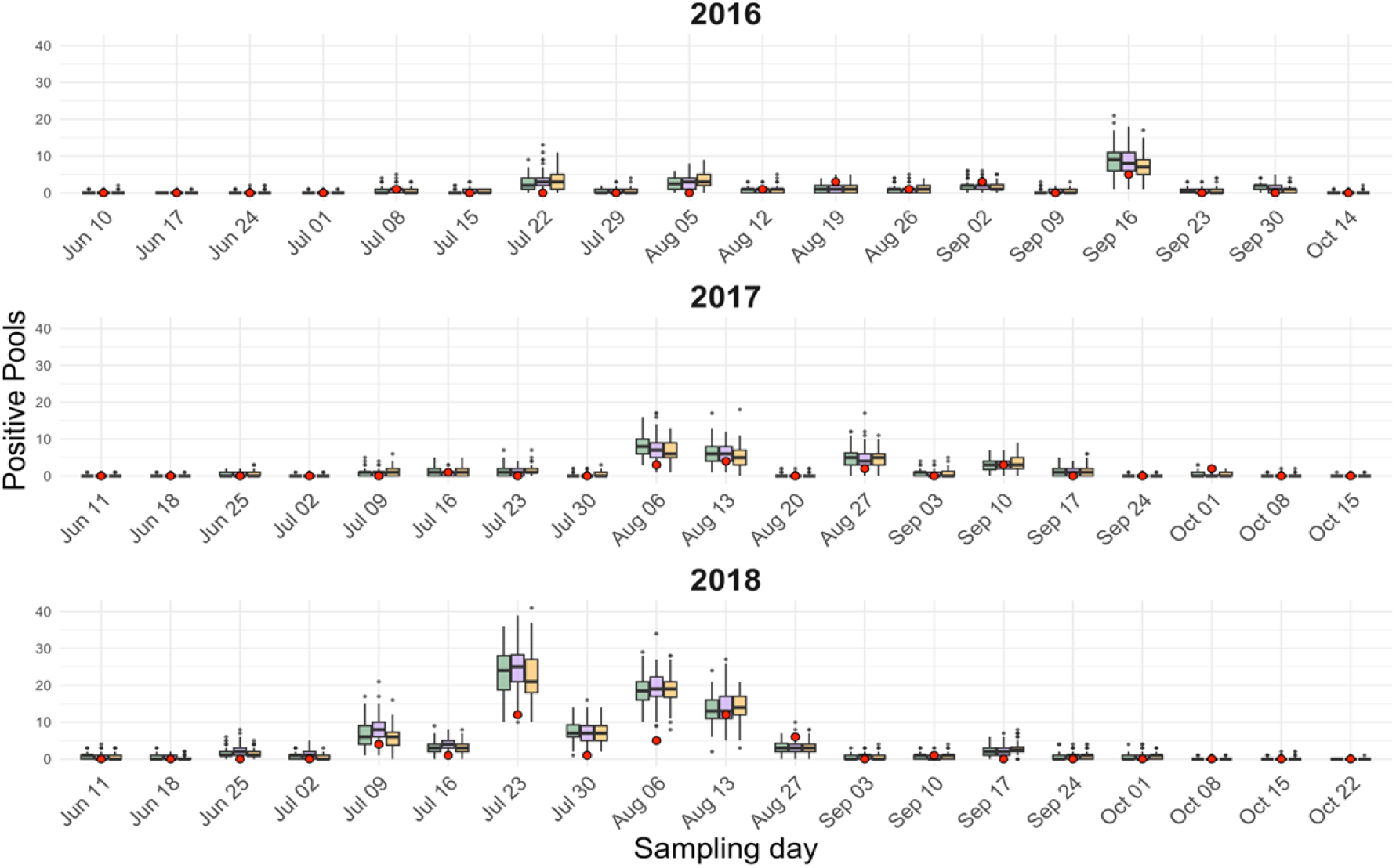
Yearly fit of the model simulations. Yearly fit of the model: red dots represent the observed mosquito prevalence. The boxplot displays the distribution of the predicted number of WNV-positive pools, corresponding to the three different models using the three target bird species (orange for blackbird, green for magpie and purple for mallard). Lower and upper hinges of the boxplot represent the 25^th^ and 75^th^ percentiles, the central line represents the median and the whiskers extending to the most extreme values within 1.5 times the interquartile range (IQR).

Blackbird contributed most consistently to WNV transmission across all three years, accounting for a mean minimum of 5% and a maximum of up to 86% of infected mosquitoes originating from bites on this species (7.7-57.6% in 2016; 9-31.4% in 2017; 5.3-86.3% in 2018). They are followed by magpie (1-1.3% in 2016; 1.2-5.7% in 2017; 1.1-33.4% in 2018), while mallard contribute less than 1% of infections (0.003-0.02% in 2016; 0.007-0.3% in 2017; 0.003-0.3% in 2018). All three species show increased contributions later in the season, with the highest contributions observed in 2018 (Fig 9).

**Fig 9.**
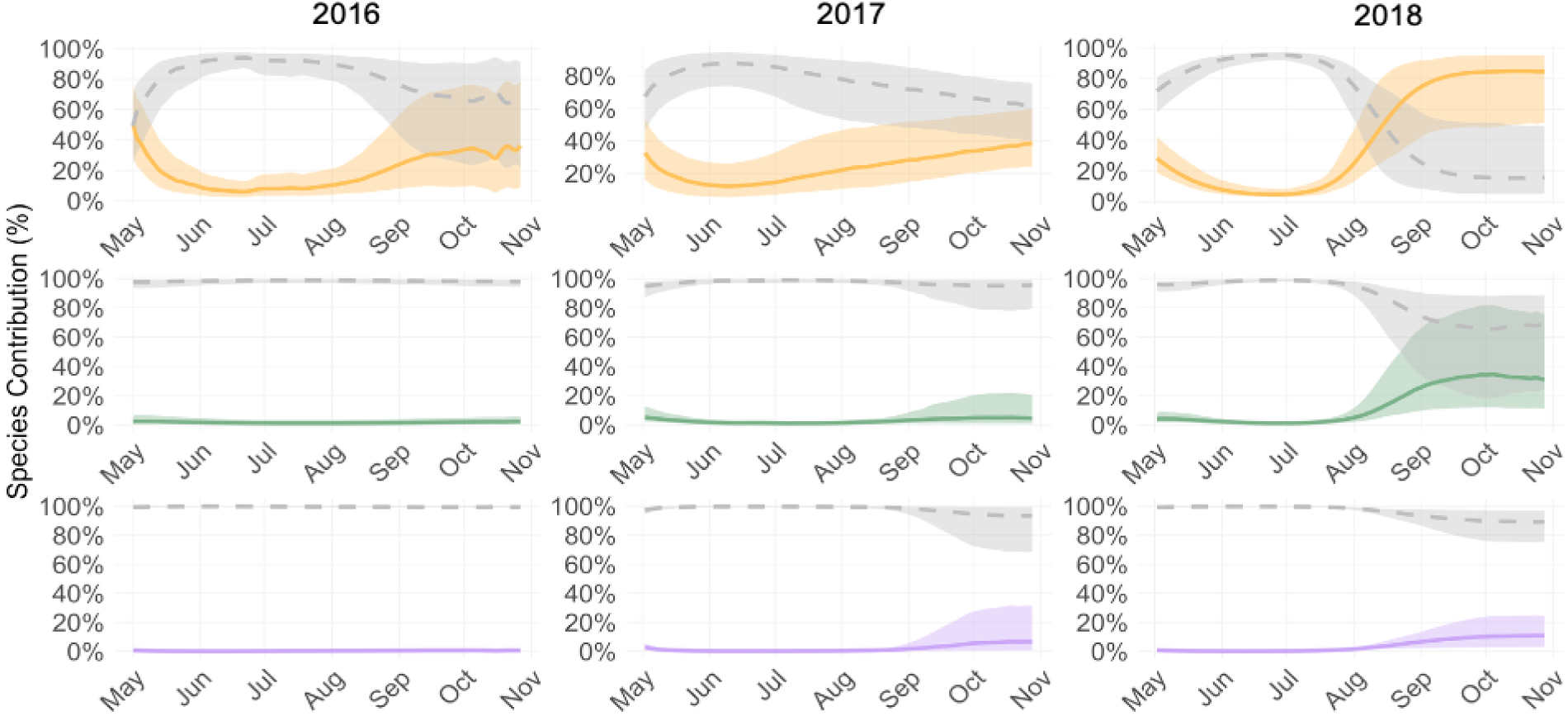
Species contribution to WNV dynamics. Daily contribution to new WNV-infections by blackbird (orange, top row), magpie (green, middle row), and mallard (purple, bottom row) during the three years of simulation, from April to the end of October. Solid orange, green and purple lines represent the mean contribution across simulations, with shaded areas indicating the 95% confidence intervals. The dashed grey lines in the panels represent the mean contribution of the remaining avian competent community to WNV spread, with its 95% CI.

## Discussion

We applied mathematical modelling to examine the influence of avian demographic traits, such as the number of fledglings (driven by the birth rate *µ_B_*), hatching season synchrony (*s*), and timing of hatching (*φ*), on WNV dynamics. The analysis demonstrates that all investigated avian demographic factors significantly affect the daily number of infectious mosquitoes. In particular, the timing of hatching affects WNV dynamics increasing the predicted number of infectious mosquitoes as the birth season shifts toward September. The number of newborns influences both the timing and size of the peak, while birth season synchrony primarily affects its size, though to a lesser extent. Additionally, among the simulated species, blackbird is identified as key contributors to WNV spread, whereas mallard exhibit only a marginal role.

In addition to species-specific epidemiological parameters, which directly influence infection dynamics and each species’ contribution to transmission, demographic parameters are also known to play a key role [18–21,45]. Particularly in multispecies infections such as WNV, the contribution of each species to disease dynamics arises from the intricate interplay between ecological and epidemiological factors driving infection.

WNV recurrence presents high variability in time and between areas, with unexpected peaks in the number of infectious mosquitoes in some year and area [30,41], but the drivers of this variability remain unclear, raising questions about whether it is mainly attributable to changes in climatic and environmental conditions and mosquito abundance, or whether more intricate ecological interactions between competent host and vector communities play a greater role. Differences in the epidemiological characteristics of birds have been explored in laboratory studies [8–10], and their relevance to WNV dynamics has been demonstrated [28]. However, little is still known about how demographic traits of birds influence WNV dynamics. Field investigations remain challenging and demanding due to the large number of species involved and the complexity of conducting experiments in natural settings [26,46]. In this context, a theoretical investigation like the one we propose can help in complementing field studies, enhancing their results and highlighting both strengths and weaknesses in current knowledge. In fact, by examining which and how the demographic characteristics of the host population influence WNV transmission, our analysis provides a framework for theoretically understanding the factors that determine WNV dynamics, but also helps to identify in practice the species whose demographic characteristics are most favourable to virus transmission. We demonstrated that a seasonal shift in the number of newborns leads to a higher WNV prevalence in mosquitoes. Additionally, a greater fledgling number results in an earlier epidemic peak. These findings constitute the principal contributions of our research, highlighting the substantial impact that even small differences in the often overlooked demographic traits of reservoir species can have, further underscoring the importance of accurately identifying reservoir species to predict the size and timing of the epidemic peak. Also, by showing that species with later hatching timing of breeding contribute more significantly to WNV spread, they provide a framework for identifying the most important avian host species by linking key demographic traits to transmission dynamics. At the same time, inter-annual differences in population dynamics of avian species may contribute to the difference observed in WNV prevalence between years. These findings can guide field and laboratory studies toward the most relevant species while also highlighting the gaps and uncertainties in our understanding of the disease. Furthermore, given that human cases typically emerge by the end of the summer season (August/September, [47]), our results highlight the importance of identifying species with low reproductive output, as these may delay the peak of WNV prevalence towards September, thereby potentially increasing the risk of human infection. It should also be noted, however, that the model does not explicitly account for age groups; further refinement of the modelling framework to include age groups could provide a better understanding of the role of young birds in the dynamics of WNV.

In the second part of our study, of the three representative species, we identified blackbird as a key species in the seasonal dynamics of WNV. Their offspring production falls between that of magpie and mallard, with a birth peak occurring later than mallard but earlier than magpie, but they also experience higher mosquito biting rates than other species [43]. Despite blackbird is here intended to represent a group of species sharing similar demographic traits, to ensure realism, species-specific epidemiological parameters were used. Worth noting that mosquito biting rate toward blackbird is higher if compared to the one of other species included. The results obtained therefore do not conclude that blackbird are responsible for the circulation of WNV. Instead, it demonstrates that a combination of species-specific traits, including demographic traits, determines a species’ potential role in infection dynamics, and that bird species beyond the traditionally studied one may contribute to transmission. This finding also aligns with previous works highlighting the critical role of mosquito biting rates in WNV dynamics [28,43]. However, while further investigations are needed to elucidate the specific role of blackbird in WNV transmission; our results suggest that given their demographic and epidemiological characteristics, this species (along with potentially other species exhibiting similar demographic traits) could play a pivotal role in WNV dynamics. Consequently, these findings underscore the critical importance of avian demographic characteristics in shaping WNV transmission patterns and indicate that variability in disease spread may be contingent upon shifts in bird population dynamics.

Therefore, in addition to the mechanisms traditionally considered to drive WNV (like temperature and bird immunity), the interannual differences observed in the contribution of different species may suggest that local ecological conditions significantly influence WNV transmission. For instance, the higher circulation observed in 2018 compared to 2016 and 2017 prompts consideration of whether changes in the dynamics, abundance or relative abundance of specific bird populations, might have played a role in amplifying virus transmission that year.

These findings reinforce the need to complement theoretical approaches with field-based studies conducted under more controlled conditions, to further assess and validate the specific role of the blackbird in WNV dynamics. On top of that, this analysis of the individual contributions of host species ultimately emphasizes the need to consider both epidemiological and demographic characteristics of bird species in understanding WNV dynamics.

Our analysis also revealed that the contribution of all investigated species to WNV spread increases from July/August onward, reflecting, with a slight delay, the rise in susceptible individuals following births (hatching). This finding emphasizes the relationship between the timing of the bird birth and the subsequent increase in virus transmission, further underscoring the importance of accounting for demographic traits in epidemiological investigations.

A limitation of the present work is its reliance on broad-scale estimates for mosquito abundance and avian traits, which may not adequately represent local conditions affecting WNV transmission. Given these spatial constraints and the approximations required for parameterizing demographic functions, future studies incorporating fine-resolution, location-specific avian population dynamics would substantially enhance the ecological realism of our findings. Additionally, the models were parameterized using only three years of observational data; the incorporation of a longer temporal dataset would likely enhance the robustness of model estimates and the ecological realism of simulation outcomes. Also, our model does not explicitly calibrate parameters on bird prevalence and immunity, thus overlooking potential effects on WNV dynamics of changes birds immunity between years [48]. Nevertheless, this work establishes a solid and novel foundation for future research and emphasizes that demographic characteristics remain an essential factor that must not be disregarded in WNV transmission studies. By highlighting the importance of demographic and ecological characteristics of host species in pathogen spread, we showed that a joint contribution of ornithologists and ecologists is crucial in the context of WNV, as knowledge in species abundance, ecological traits, and regional differences provides essential data to refine and enhance epidemiological investigations. Effective communication between disciplines such as epidemiology and ecology is fundamental and there is a growing need for multidisciplinary frameworks to study the spread of infectious diseases [49–51].

At last, comparing these new results with our previous findings [28], we conclude that incorporating a continuous birth function improves the representation of bird population dynamics by considering continuous changes in birth rates over time, despite the necessary approximations involved in translating demographic characteristics into parameters, and aligns more closely with natural reproductive patterns. While improvements to the accuracy of the functions are feasible, these improvements fell outside the exploratory scope of the present paper and its focus on WNV dynamics. While our model is based on data from a specific region and time frame (Lombardy, 2016–2018), its structure and framework ensure flexibility and generalizability, making it a robust tool for understanding and predicting WNV dynamics in different contexts. Despite that, our simulations were based on a closed bird population, excluding the effects of migratory processes, emigration and immigration, which future studies could incorporate. Additionally, modelling choices can influence results [52], and the combined effect of parameters may also impact pathogen spread [28].

In conclusion, our results indicate that avian demographic traits, particularly the timing of reproduction, can shape WNV transmission dynamics, with later birth seasons associated with amplified epidemic peaks, and variation in the number of newborns influencing both the timing and magnitude of these peaks. When demographic and epidemiological traits are jointly considered in the interpretation of transmission patterns, species rather than corvids may emerge as contributing to WNV spread, and blackbird, given its abundance and ecological/epidemiological traits, might be one of the key species on which to focus WNV surveillance efforts in the study region. These findings are intended to inform mechanistic understanding and surveillance prioritisation, and to underscore the need for close interdisciplinary collaboration to improve our understanding of host–vector systems and support integrated surveillance strategies, rather than to motivate interventions targeting specific wildlife species.

## Acknowledgements

Work partially supported by UNIMI GSA-IDEA Project.

## Author Contributions

Conceptualization, E.F. and N.F.; methodology, E.F., G.M., N.F.; formal analysis, E.F. and E.C.; investigation, E.F. and E.C.; data curation, E.C, E.F., L.I., D.L., M.P.C. and G.M; writing—original draft preparation, E.F.; all authors contributed to writing—review the ms; visualization, E.F. and E.C.; supervision, N.F.; project administration, N.F.; funding acquisition, N.F. All authors have read and agreed to the published version of the manuscript.

## Supporting information captions

**S1 Text.** Birds’ demographic parameters

**S2 Text.** Description of base model equation and parameters

**S3 Table. Data table.** Recorded average entomological captures and number of positive pools for each year.

